# Plant host identity drives *Andropogon gerardii* rhizobiome assembly strategies under increasing abiotic stress

**DOI:** 10.1101/2025.04.12.648431

**Authors:** Brooke Vogt, Anna Kazarina, Jack Sytsma, Bryttan Adams, Leslie Rodela, Dasik Clouse, Reece Keller, Darcy Thompson, Helen Winters, Maggie Wagner, Ari Jumpponen, Loretta Johnson, Sonny T.M. Lee

**Affiliations:** Division of Biology, Kansas State University, Manhattan, Kansas, USA; Department of Ecology and Evolutionary Biology and the Kansas Biological Survey, University of Kansas, Lawrence, Kansas, USA

**Keywords:** Microbiome, Microbial Community Assembly, Host-microbe Interactions, Climate Change, 100^th^ Meridian, Plant-microbe interactions

## Abstract

Predicted changes in precipitation threaten tallgrass prairies by altering the soil microbial communities that are essential for plant resilience. *Andropogon gerardii*, a dominant grass in tallgrass prairies, spans the contiguous North American precipitation gradient. However, it remains unclear to what extent the rhizosphere microbiomes (rhizobiomes) are influenced by the plant-host environmental interaction. To assess how environmental and host factors shape the rhizobiome, we surveyed *A. gerardii* populations across 25 remnant prairie sites (June–August 2023) within its native range in the United States, characterizing the microbiomes in the rhizosphere and soils using 16S amplicon sequencing. We demonstrated that while geographic location largely structured both rhizosphere and soil communities, regional precipitation (60-day rainfall) emerged as a primary driver of the microbial community assembly. We observed distinct microbial divides across the dry and wet regions of the North American “arid-humid divide.” Importantly, we found the first compelling large-scale evidence that regional precipitation has a profound influence on rhizobiome assembly. In the most arid regions, rhizosphere microbial communities exhibited significantly more predicted stochasticity than those in the local soil and contained taxa related to host-benefiting functions. Our study suggests that intensified host-driven selection for specific microbial variants occurs under heightened abiotic stress, highlighting the host’s pivotal role in shaping its rhizobiome composition in challenging environments.

## Introduction

Changes in the assembly and function of plant host-associated microbiomes can be influenced by a mosaic of factors, such as abiotic stress induced by climate change^1–4^. Over the past 10,000 years, the dominant tallgrass prairie grass, *Andropogon gerardii,* has evolved into genetically distinct ecotypic varieties based on precipitation (dry, mesic, and wet ecotypic varieties) across North America^5–9^. Microevolutionary processes have modified the morphological and physiological traits of these varieties, enabling them to survive and thrive in regions with varying precipitation levels, temperatures, and soil physicochemical properties^6,10–12^.

Some studies suggest that plant host-associated rhizobiomes are specific to the plants’ locations, indicating that plants of the same host variety and across different host varieties are distinct^13,14^. This view asserts that rhizobiome assembly is more strongly influenced by various local abiotic stresses as well as environmental and edaphic characteristics than by regional similarities in precipitation and climate. Other studies propose that rhizobiomes of the same host variety are similar across various locations, whereas those of different host varieties differ from each other^15,16^. According to this perspective, precipitation-based abiotic stress drives the host plants to select similar microorganisms, resulting in analogous rhizobiome communities within the same host variety^17,18^. These contradicting hypotheses suggest a lack of detailed information on how abiotic, precipitation-based stress affects the rhizobiomes of *A. gerardii* varieties.

Prior reciprocal garden experiments^19,20^ examining the rhizobiomes of the dry, mesic, and wet varieties of *A. gerardii* have supported the idea that host-associated rhizobiomes are primarily influenced by regional precipitation. In these studies, *A. gerardii* recruited rhizobiomes similar to those at its native sites, supporting the home-field advantage hypothesis^19,20^. If microbial assembly is driven by drought stress, then different forms of this stress should lead to the host selecting for different specialists.

While the differences among dry, mesic, and wet varieties of *A. gerardii* have evolved over thousands of years due to natural selection, how quickly their host-associated rhizobiome can adapt to long-term changes in precipitation remains to be seen. The consequences of a rhizbiome’s ability to adjust to different forms of precipitation-based abiotic stress are crucial. In this century, the 100^th^ meridian (“arid humid” divide) has shifted eastward, a trend predicted to accelerate due to the compounding effects of climate change^5,21–24^. Understanding the host rhizobiome assembly and its vulnerabilities is crucial, mainly because *A. gerardii* constitutes up to 80% of the biomass in tallgrass prairies^25^. In addition, central tallgrass prairies are critically endangered, with only 4.4% remaining in North America^6,8,25–28^. Rapid shifts in precipitation patterns may lead to a decline in *A. gerardii* if their microbiomes are closely linked to local ecological and soil microbial communities, and cannot adapt to long-term precipitation shifts. If *A. gerardii* microbiomes are solely influenced by local conditions, then long-term variations in rainfall could pose a significant threat to the host’s health, longevity, and survival.

In this current study, we aimed to evaluate how regional precipitation conditions influence rhizobiome assembly. We further investigated whether the plant host variety has a greater influence on the associated rhizobiomes in areas experiencing more significant precipitation-based abiotic stress. To understand the composition of the microbial communities associated with the ecotypic varieties of *A. gerardii*, we conducted a biogeographical survey of 25 locations across the contiguous United States. We analyzed 16S rRNA amplicon data from local soil (LS) and plant host-associated rhizobiome (RB) samples. Our study addressed three key questions: I) How do local and regional characteristics influence microbial community composition? II) How does plant host selection affect rhizobiome composition and diversity compared to local soil, and is biotic filtering consistent across abiotic stress levels? III) How do local soil and rhizobiome communities compare in similarity across different precipitation-related host variety ecotypes?

## Methods

### Sample collection and preparation

Sample collection took place between June and August of 2023. We selected 25 sample locations based on the historical distribution of *A. gerardii*, aiming to represent variations in precipitation^29^, soil physicochemical properties, climate, and host variety across the contiguous United States (Supplementary Table S1). We collected local soil and rhizosphere samples from six haphazardly chosen *A. gerardii* plants at each location. From each plant, we sampled one soil core (10 cm deep and 1.25 cm in diameter) near the base of the plant tillers. This core sample was manually homogenized in a sterile plastic bag. From the homogenized sample, three aliquots of roots (approximately 20 cm in length each) were manually collected, gently shaken to remove any loose soil, and placed into three microcentrifuge tubes. From the remaining soil, we collected three soil aliquots (∼1g each) into three microcentrifuge tubes.

Any remaining soil (without roots) from the six plant replicates that were not used for the local soil or rhizobiome samples were stored in sterile bags on ice for the soil physicochemical property analyses (n=25 samples). All samples were stored on ice in a cooler until they were frozen at -80°C within 96 hours after sampling for further processing.

### Soil physicochemical properties

The soil physicochemical property analyses were conducted at Kansas State University’s Soil Testing Laboratory (https://www.agronomy.k-state.edu/ outreach-and-services/soil-testing-lab/soil-analysis/researchers.html). The analyses were performed on the local soil samples (n = 25). Detailed procedures are in the revised version of the Recommended Chemical Soil Test Procedures for the North Central Region manual^30,31^. However, a summary of the methods for each test is provided below.

Samples were analyzed for various physicochemical properties, including soil pH, total nitrogen, carbon content (both organic and inorganic), phosphorus content, and texture (percentages of sand, silt, and clay). Before analyses, the samples were dried in a drying oven at 60 °C overnight. The dry soils were passed through a 2 mm sieve to break up aggregates and ensure homogeneity.

For pH analysis, soil was mixed with an equal amount of deionized water in a 1:1 solution^30,32–34^. The mixture was stirred for 5 seconds or until a uniform soil-water slurry was achieved. After allowing it to sit for 10 minutes, pH measurements were taken using a Skalar SP50 Robotic Analyzer (Skalar Inc.; Georgia, USA).

Phosphorus concentrations were measured in parts per million (ppm P) using the Mehlich III method on 2 grams of soil^30,35,36^. The soil was suspended in a 1:10 solution with the standard Mehlich III solution, which consists of 0.2 N acetic acid (CH COOH), 0.25 N ammonium nitrate (NH NO), 0.015 N ammonium fluoride (NH F), 0.013 N nitric acid (HNO), and 0.001 N EDTA (ethylenediaminetetraacetic acid)(HOOCCH_2_)_2_NCH_2_CH_2_N(CH_2_COOH)_2_). The mixture was shaken at 200 excursions per minute for 5 minutes. After shaking, the slurry was filtered through a Whatman No. 42 filter. From the clear filtered slurry, 2 mL was transferred to a test tube, and 8 mL of the working solution was added. The working solution was prepared by combining 25 mL of Murphy-Riley Solution A (which contains 0.051 N ammonium molybdate [(NH) Mo O “4H O] and 0.004 N antimony potassium tartrate [K Sb (C H O) ”3H O]) with 10 mL of Solution B (0.749 N ascorbic acid [C H O ]) in a total volume of 1 L.

After letting the mixture sit for 10 minutes, the samples were loaded in triplicate onto a 96-well plate and analyzed using a Lachat QuikChem 8500 Series 2 photometric colorimeter (Hach Inc.; Colorado, USA) at a wavelength of 882 nm, following the orthophosphate method (12-115-01-1-A). Phosphorus measurements were determined by comparing the samples against a 500 ppm potassium dihydrogen phosphate (KH PO) stock solution, diluted to a 50 ppm standard solution before loading into the 96-well plate.

Total nitrogen and total carbon (including organic and inorganic components) analyses were conducted on 0.35 grams of soil using a LECO TruSpec CN analyzer (LECO; Michigan, USA), following the procedures outlined in the operation manual^37^. The samples were encapsulated and securely sealed in the loading head. They were then purged to eliminate any atmospheric gases that could interfere with the measurements. Once all atmospheric gases were removed, the samples were combusted at 950°C to ensure rapid combustion while flushed with pure oxygen. A consistent aliquot of gases was then homogenized and separated. The total percentage of carbon and nitrogen was calculated based on weight percentages using the formula: Total % of Carbon or Nitrogen = (Mass of C or N / Total Sample Mass) * 100. The analysis showed a standard deviation of 2.18% for soil carbon and 3.35% for soil nitrogen.

Soil texture was determined using the modified hydrometer method^38,39^ on a 50-gram soil sample. First, the soil was placed in a dispersion cup, and then 100 mL of a 5% sodium hexametaphosphate solution (Na_6_(PO_3_)_6_) was added. The mixture was stirred for 30 to 60 seconds until uniform. This slurry was then transferred to a 1000 mL cylinder filled to the 1000 mL mark with deionized water. The mixture was allowed to sit overnight to settle. To begin the experiment, a plunger was inserted, and the sample was mixed for an additional 30 seconds to ensure homogeneity. A Bouyoucos hydrometer (Fisherbrand™; Soil Analysis ASTM Hydrometer) was then placed in the slurry. After 40 seconds, the initial hydrometer reading was recorded. A second reading was taken after 2 hours.

The percentages of sand, silt, and clay were calculated using the following formulas: - % Clay = (Final Reading * 100 / mass of sample) - % Silt = (Initial Reading * 100 / mass of sample) - % Clay - % Sand = 100% - % Silt - % Clay All readings were adjusted using a blank sample (100 mL of the 5% dispersion solution plus 880 mL of deionized water) and applying temperature and density corrections. For every one °F above 68 °F, 0.2 was added to the reading, while for every one °F below 68 °F, 0.2 was subtracted from the reading.

### DNA extraction, 16S rRNA amplicon sequencing and analyses

We extracted DNA from the samples using the E.Z.N.A.^®^ Soil DNA Kit (Omega BIO-TEK; Georgia, USA) following the manufacturer’s protocol with minimal modifications^40^. The only adjustment for the local soil samples was the use of glass beads for cell lysis, performed with a Qiagen TissueLyser II (Qiagen; Hilden, Germany) at 30 revolutions per second for 10 minutes. For each of the 25 locations, we extracted three aliquots separately and selected one from each of the six plants based on DNA concentration and quality. The final elution volume for each local soil sample (n = 150) was 100 μL.

Similar to the soil samples, we extracted DNA from the rhizosphere samples using the E.Z.N.A.^®^ Soil DNA Kit (Omega BIO-TEK; Georgia, USA) but with significant modifications to the manufacturer’s protocol to ensure optimal DNA quality and yield. For these samples, we combined three aliquots into one during the binding phase of the protocol. Additionally, we used liquid nitrogen to physically macerate the roots into a fine powder, and then lysed the tissues using the TissueLyser II for 16 minutes at 30 revolutions per second. Depending on the sample location, we adjusted the final elution volumes for the rhizobiome samples, ranging from 30 μL to 150 μL.

We used the Qubit™ dsDNA Broad-range (BR) DNA quantification kit on a Qubit™ 4 Fluorometer (Thermo Fisher Scientific, Invitrogen™; Massachusetts, USA) to determine the sample eDNA concentration. The quality of the DNA was assessed by measuring the A260/A280 and A260/A230 ratios on a Nanodrop™ One Spectrophotometer (Thermo Fisher Scientific, Invitrogen™; Massachusetts, USA). After quantifying the DNA, we normalized each sample to a concentration of 5 ng/μL using molecular-grade DNA/RNA-free water.

The normalized eDNA was sequenced at the Kansas State University Integrative Genomics Facility (https://www.k-state.edu/igenomics/). The DNA amplification was performed using the 515F and 806R primers^41^ (16S rRNA v4 amplicon region), and sequenced with 2 x 300 cycles on the Illumina MiSeq (Illumina; California, USA). We utilized QIIME 2 v. 2024.5^42^ to process the raw reads. We applied the QIIME 2 cutadapt^43^ plugin to remove the primer sequences.

Subsequently, we eliminated sequences with an error rate greater than 0.1 and those with undetermined bases or without a primer. For further quality control, we used DADA2^44^ with consistent parameters across various runs to trim the reads, ensuring that the 25^th^ percentile of reads had a quality score above 15 (Forward = 270 and Reverse = 250). To mitigate biases caused by differences in sequence depth across samples in subsequent diversity analyses, we rarefied the dataset to 15,000 reads per sample. We used the QIIME 2 pipeline to compute alpha (Shannon’s Entropy (H’), Faith’s phylogenetic diversity (Faith PD), and number of observed features (S_obs_)) and beta diversity metrics. Finally, we employed the pre-trained classifier in QIIME 2 for bacterial taxonomic classification, referencing the SILVA v. 138 database^45^. After generating the ASV table, we removed all ASVs assigned to Eukarya, chloroplasts, or mitochondria, as well as those that remained unresolved or unclassified at the domain level, or included non-specific reads before statistical analyses.

To methodically tease apart the influence of environment and plant host on the rhizobiome composition, we analyzed beta dispersion to examine how community assembly strategies influenced the distance to the centroid between dry and wet local soil and rhizobiome samples, focusing on whether sample distribution was driven more by stochastic (random) or deterministic processes. We defined samples where beta dispersion distances close to 1 indicate increased stochasticity, whereas smaller distances (0.8) demonstrate more predictability^46^.

To assess specific microbial families that contribute to stochasticity, we performed oligotyping analysis. For this analysis, we employed the QIIME 2 oligotype function^47^ (v. 3.2) and the minimum entropy decomposition (MED) algorithm^48^ to identify oligotypes within our aligned 16S rRNA amplicon sequences. We aligned our sequences using the MUSCLE function^49^. To ensure an unbiased taxonomic assessment of oligotypes across various sample types, we set the number of components to 7 with a minimum substantive abundance parameter of 1 and included all features. These parameters were determined based on the entropy analysis of oligotype entropy peaks. We retained (LS: n = 2,248) and (RB: n = 2,876) oligotypes in this analysis.

In our taxonomic evaluation, we extracted specific features belonging to specific families from our previous taxonomic assessment (SILVA v. 138^45^). We filtered features that belonged to the *Micromonosporaceae, Nitrosphaeraceae*, *Xiphinematobacteraceae, Vicinamibacteraceae, Rhodanobacteraceae,* and *Pedosphaeraceae* families because of their significant representation and overall abundance across all samples. We set the number of components to 2 for the taxonomic analysis while maintaining a minimum substantive abundance parameter of 1. Following the taxonomic oligotyping analysis, *Micromonosporaceae, Nitrosphaeraceae*, *Xiphinematobacteraceae, Vicinamibacteraceae, Rhodanobacteraceae,* and *Pedosphaeraceae* families yielded the following total number of oligotypes: (LS: n = 9 and RB: n = 10; LS: n = 9 and RB: n = 10; LS: n = 14 and RB: n = 13; LS and RB: n = 16; LS: n = 6 and RB: n = 7; and LS and RB: n = 16).

### Statistical analyses and data visualization

We used RStudio version 4.4.2^50,51^ for all statistical analyses and figure generation. We defined significance for all tests as having a p-value or adjusted p-value (used when appropriate, given the analysis) less than 0.05.

For our studies of soil/climate properties, alpha diversity, and relationships between these factors, we utilized the car^52^, dplyr^53^, devtools^54^, dunn.test^55^, pairwiseAdonis^56^, tidyr^57^, tidyverse^58^, ggplot2^59^, FSA^60^, broom^61^, rstatix^62^, and vegan^63^ packages to evaluate how unique the soil’s physicochemical properties were based on the sample’s location, precipitation, and sample cluster category.

We used one-way ANOVAs detect differences among the factors including soil pH, phosphorus concentration (ppm P), carbon and nitrogen content (% total C and N), C:N ratio, soil texture (% sand, % silt, and % clay), and precipitation (rainfall) concerning the sample location and cluster category (dry, ambiguous, and wet). Furthermore, we analyzed the alpha diversity metrics, sample location, cluster, and pH/precipitation gradients using a one-way ANOVA to compare the means of the S_obs_, Faith’s PD, and H’ for the local soil and rhizobiome samples. We performed a two-way ANOVA to analyze the relationship between our alpha diversity metrics, cluster categories, and pH/precipitation. Additionally, we also used a one-way ANOVA to define statistical differences among the beta dispersion means for each location and sample type. While ANOVA identified differences among the factors, we performed Tukey HSD post hoc tests to determine where those differences were. Additionally, we applied the Kruskal-Wallis rank sum test to evaluate pairwise comparisons between sample locations and clusters for both the local soil and rhizobiome samples, analyzing our soil properties and precipitation (rainfall) data. Using both parametric and non-parametric approaches allowed us to validate our findings across different statistical assumptions and ensured the robustness of our results. Finally, we conducted PERMANOVA analyses to determine whether bacterial community composition varied in relation to location, precipitation cluster category, soil properties, and precipitation. We then performed PERMANOVA pairwise comparisons to explore the pairs of groups individually between the factors described in our PERMANOVA analyses. In our analysis of how microbial diversity is impacted by climate and soil properties, we performed both t-test and linear regression analyses to further explore these relationships.

To assess differences in the variability of community composition within each cluster category and evaluate category-specific dispersion patterns, we conducted analyses of multivariate homogeneity of group dispersions (employing the betadisper function). Using the car^52^ and vegan^63^ packages, we assessed the differences between groups based on the mean distance to the centroid for each category. This evaluation helped us determine the probability of overlap between the cluster categories. Within our betadisper test, we performed a PERMANOVA, a permutation test, and a pairwise permutation test to examine sample heterogeneity within each cluster category. Finally, we used Pearson’s Chi-squared tests in the tidyverse^58^ and readr^64^ packages to find statistical differences in ASV composition between sample types (LS and RB) and the cluster categories.

We used beta dispersion distances and the beta-group-significance function in QIIME 2 to infer stochasticity within a location and between sample types, thereby surmising the underlying strategies of community assembly. For the oligotyping analysis, we employed the beta dispersion tests previously described.

We utilized the RStudio packages ggplot2^59^, patchwork^65^, dplyr^53^, multcomp^66^, multcompView^67^, agricolae^68^, tidyr^57^, venn^69^, tidyverse^58^, vegan^63^, cluster^70^, sf^71^, maps^72^, reshape2^73^, pheatmap^74^, RColorBrewer^75^, car^52^, gridExtra^76^, phyloseq^77^, igraph^78^, ggnewscale^79^, readr^64^, ggthemes^80^, ggtext^81^, ggVennDiagram^82^, randomcoloR^83^, stringr^84^, and VennDiagram^85^ to visualize the analyses performed in this study. All raw sequence data for this study are publicly available in the NCBI SRA under BioProject accession number PRJNA1249957. All bioinformatic and R codes are deposited in GitHub (https://github.com/SonnyTMLee/Plant-host-identity-drives-rhizobiome-assembly-under-increasing-abiotic-stress.git).

## Results and Discussion

We processed a total of 13,087,481 (local soil) and 16,677,859 (rhizobiome) raw reads. After quality control, we retained 11,759,436 reads (2,462 ASVs) for the local soil and 14,627,083 reads (3,039 ASVs) for the rhizobiome (Supplementary Table S2). In the local soil samples, 92% (2,267 ASVs) of the ASVs resolved to the genus level, and 64% (1,567 ASVs) resolved to the species level. We observed similar resolution across the rhizobiome samples, where 93% (2,815 ASVs) resolved to the genus and 65% (1,985 ASVs) resolved to the species level. The five most abundant phyla in the local soil samples were Proteobacteria (22%, 532 ASVs), Actinobacteriota (11%, 278 ASVs), Bacteroidota (9.8%, 242 ASVs), Chloroflexi (8.1%, 200 ASVs), and Acidobacteriota (7.6%, 187 ASVs) (Supplementary Table S3). We observed similar dominant phyla across the rhizobiome samples, with Proteobacteria (23%, 691 ASVs) being the most dominant, followed by Actinobacteriota (10%, 315 ASVs), Bacteroidota (9.9%, 302 ASVs), Planctomycetota (7.5%, 229 ASVs), and Chloroflexi (7.5%, 228 ASVs) (Supplementary Table S3). Of all our samples, 0.12% (3 ASVs) from the local soil and 0.13% (4 ASVs) from the rhizobiome were unclassified.

### Regional precipitation shaped both local soil and plant host-associated microbial communities

We used non-metric multidimensional scaling (NMDS) and optimal silhouette scores from k-means clustering, and identified two distinct clusters among both the local soil (silhouette score = 0.21) and rhizobiome (silhouette score = 0.22) samples (Figure 1A, 1B). We observed that regional precipitation significantly influenced the two identified clusters - “wet-categorized” and “dry-categorized”. Samples in the transition zone (overlap) between the wet and dry were classified as “ambiguously-categorized.” The ten samples organized as dry-categorized local soil (DC_LS_) were: MT1, SD1, ND1, MN1, CO1, KS2, NM1, NE2, OK1, and TX3. Three samples (KS1, NE1, and IA1) belonged to the ambiguously categorized local soil (AC_LS_), and 12 samples (WI1, MI1, IN1, MO1, IL1, TX1, TX2, LA1, AL1, NC1, and SC1) were organized as wet-categorized local soil (WC_LS_). The dry-categorized local soil samples had the same assignments as the dry-categorized rhizobiome samples (DC_RB_). However, the sample assignments exhibited slight changes from the local soil for the ambiguously- (AC_RB_) and wet- (WC_RB_) categorized rhizobiomes, in which IA1 and NE1 were reclassified into the WC_RB_ category. We surmised that the dry-categorized samples had more stable sample assignments than the wet or ambiguous-categorized samples due to long-lasting relationships between the dry variety of *A. gerardii* and its associated rhizobiome^6,19,86^. In contrast, the wet and ambiguously categorized clusters may be less tied to their respective host varieties, possibly due to the shifting of historical precipitation boundaries, bringing aridity to historically moist locations for centuries^24,87,88^. This change was most striking as a result of the acceleration of eastward shifts in regional precipitation zones encompassing the 100^th^ meridian, with predictive models indicating that these transitions are likely to accelerate within this century^24^.

**Figure 1:**
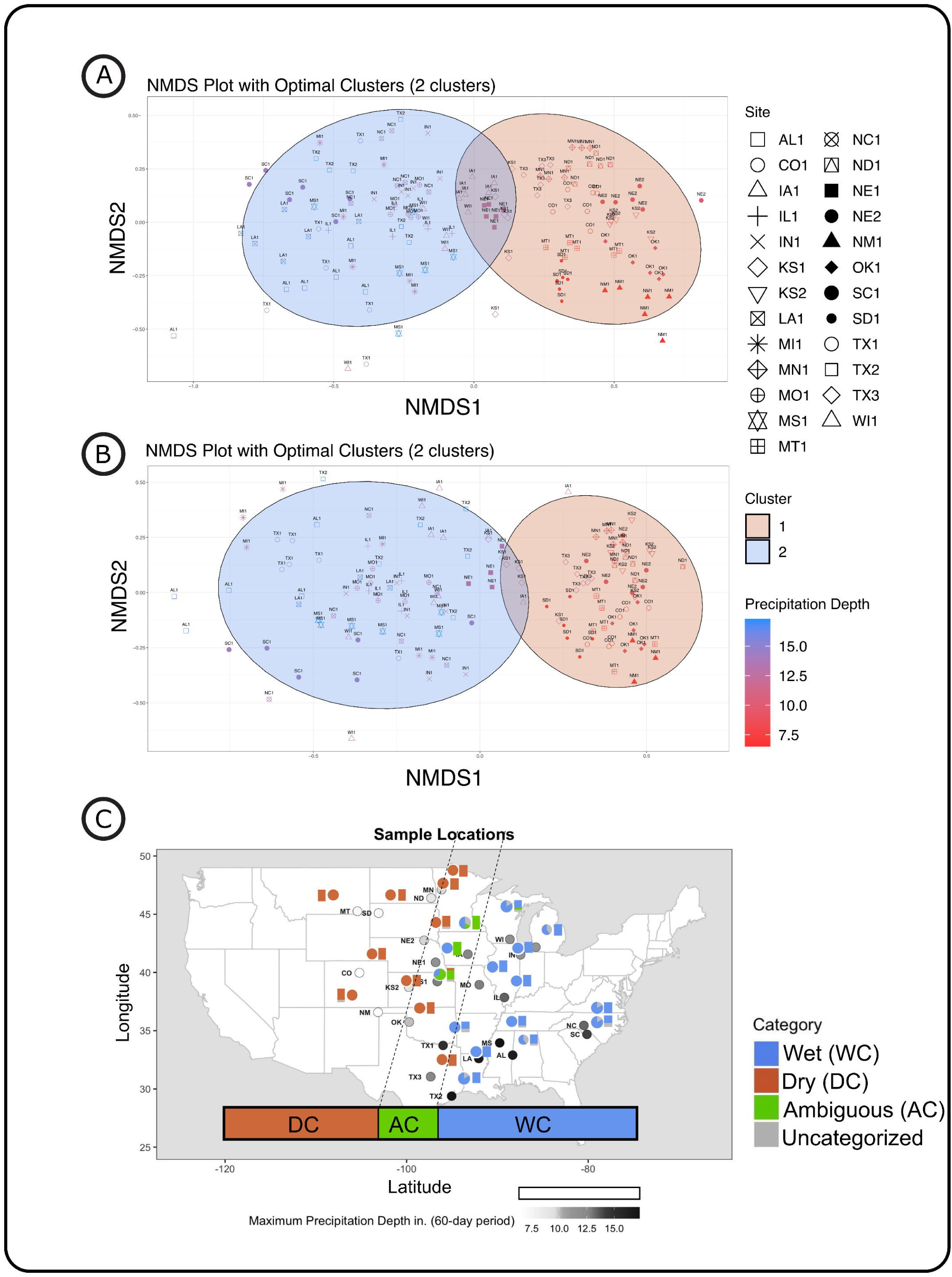
Precipitation-associated sample clustering and geospatial analysis. (A) NMDS clustering of local soil samples with k-means clustering and optimized silhouette scores showing distinct groupings with 95% confidence ellipses. Each point represents a sample, colored according to NOAA’s 60-day maximum precipitation data at the collection site. (B) NMDS clustering of rhizobiome samples with the same clustering criteria. Sample points are uniquely marked and colored by precipitation (rainfall), consistent with part A. (C) Map of biogeographical survey sites, with markers colored by NOAA’s 60-day maximum precipitation (circle) values. Stacked bar graphs within the map display the proportion of local soil samples assigned to each NMDS-defined group at each site. The adjacent pie charts summarize group assignments for rhizobiome samples.

To provide deeper insights into the impact of regional precipitation on the local soil and rhizobiome composition, we used the NOAA^29^ (v. 2) database for precipitation (rainfall) data, selected the maximum precipitation over 60 days, relative to the annual rainfall at the sample locations, and mapped the distribution of our clustering analysis across the precipitation gradient (Figure 1C). We observed a clear divide between the dry and wet categories, separating distinctly along the 100^th^ meridian, often referred to as the “arid-humid divide,” with the ambiguous samples comprising the mesic transition zone. The clustering of our samples across the 100^th^ meridian suggests that precipitation influences the microbiome composition across geographically diverse locations^89^. This indicates that the selective pressures exerted by precipitation may surpass the effects of local climate, geographic, and soil property differences^90,91^.

Our one-way ANOVA showed differences among the categories and their precipitation depth (LS: p < 2.2 * 10^-16^ and RB: p < 2.2 * 10^-16^; Precipitation: DC_LS_ = 221.5 ± 46.2 mm and DC_RB_ = 224.3 ± 45.7 mm, AC_LS_ = 312.4 ± 15.0 mm and AC_RB_ = 325.1 ± 0 mm, and WC_LS_ = 360.9 ± 55.9 mm, and WC_RB_ = 350.5 ± 54.864 mm).

Kruscal-Wallis post hoc pairwise tests showed that for the local soil, the precipitation depth between the DC_LS_ and the WC_LS_ was most significant (p-adj = 3.92 × 10^-20^), followed by the DC_LS_ vs. AC_LS_ (p-adj = 3.34 × 10^-5^) and WC_LS_ vs. AC_LS_ (p-adj = 1.02 × 10^-1^). This trend was also observed for the rhizobiome samples DC_RB_ vs. WC_RB_ (p-adj = 2.08 × 10^-9^), DC_RB_ vs. MC_RB_ (p-adj = 4.08 × 10^-2^), and WC_RB_ vs. AC_RB_ (p-adj = 1.00).

We further used PERMANOVA and showed a significant impact of rainfall on local soil and rhizobiome community structures (p = 0.001). However, though PERMANOVA pairwise analysis between our categories showed significantly different bacterial composition between all of the local soil category comparisons (adj p-values = 0.003), comparisons between WC_RB_ and AC_RB,_ as well as DC_RB_ and AC_RB,_ were insignificant (adj p-values = 0.16 and 0.07, respectively). Our results corroborated with those of another study^20^, showing that regional precipitation influences local soil communities more than plant host-associated rhizobiome communities. Here, we demonstrated the influence and regulation of the plant host on its associated rhizobiome community structure in response to adverse hydrologic conditions^1,92,93^. Next, we wanted to ask if and to what extent physicochemical soil properties contributed to this variation.

### Soil physicochemical properties and bacterial diversity were primarily influenced by geographic location rather than regional precipitation

We observed that the soil physicochemical properties (Supplementary Table S4) were more closely related to individual locations than when we performed analysis based on grouping the locations by our cluster categories: wet, ambiguous, and dry (Figure 2A). All soil properties were statistically significant between the locations (Supplementary Table S5). Nevertheless, only soil pH (LS: p = 1.05 × 10^-12^), sand content (LS: p = 1.02 × 10^-2^), and silt content (LS: p = 1.90 × 10^-4^) were significantly different across our cluster categories. We postulate that the lack of significant differences within our community-defined clusters arose because the extreme variability of individual local soil properties in specific locations was mitigated, resulting in comparable soil characteristics over extensive regions^94^.

**Figure 2:**
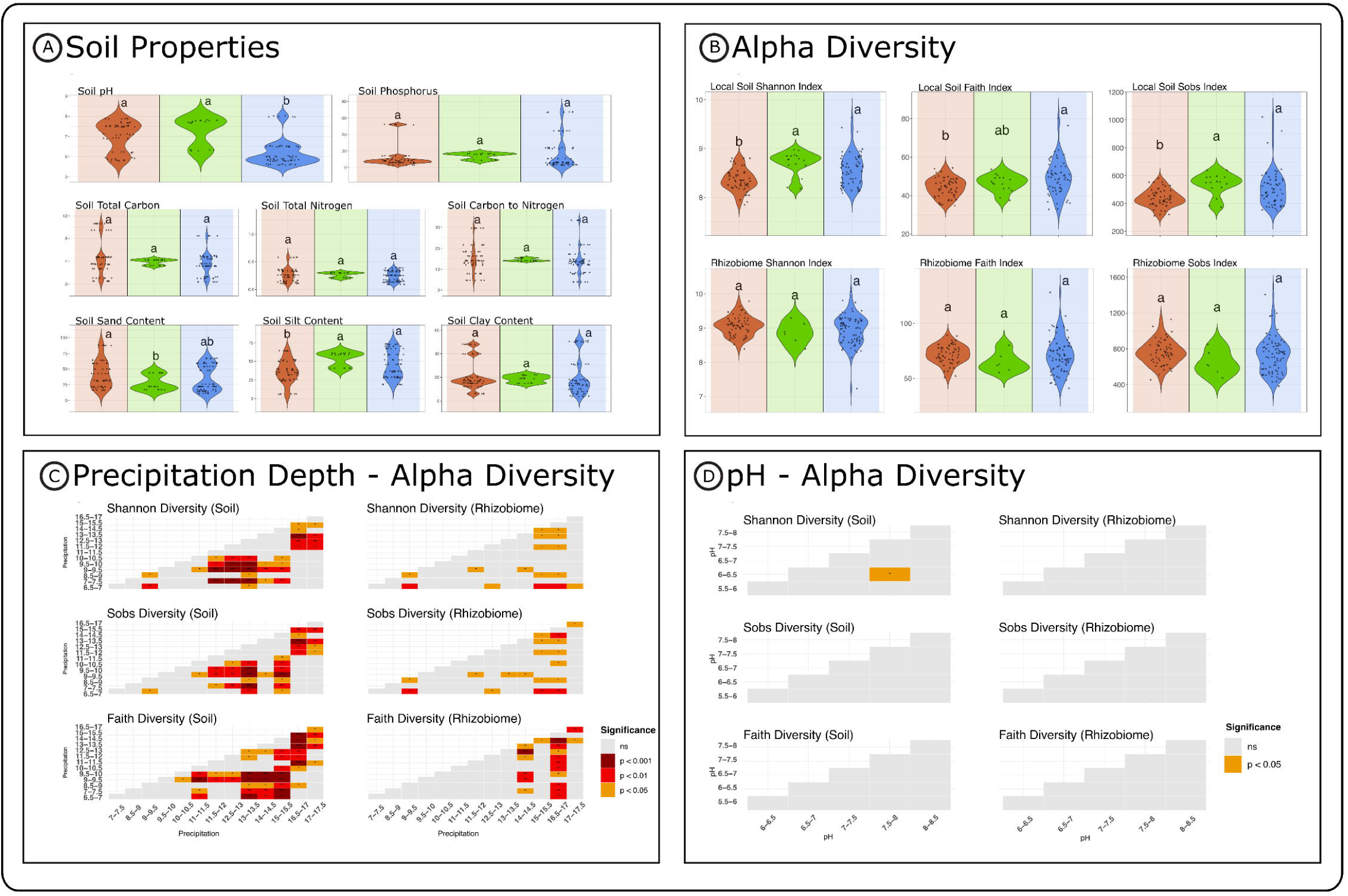
Cluster soil physicochemical properties and alpha diversity across cluster categories and soil property gradients. (A) Violin plots showing the distribution of soil physicochemical properties across assigned local soil cluster categories. Statistical differences among cluster categories were assessed using ANOVA with Tukey’s HSD post hoc tests. Red and blue panels represent dry and wet cluster categories, respectively. (B) Alpha diversity metrics, including observed features, Shannon entropy, and Faith’s PD, are grouped by cluster categories for local soil and rhizobiome samples. Significance was evaluated in the same manner as in part A. Red and blue panels represent dry and wet clusters, respectively. (C) Comparison of alpha diversity metrics between local soil and rhizobiome samples across environmental precipitation (rainfall) categories (all samples used) (assessed using independent t-tests. Statistical significance is indicated as follows: ns (grey) = p > 0.05, * (orange) p < 0.05, ** (red) p < 0.01, *** (brown) p < 0.001. (D) Comparison of alpha diversity metrics between local soil and rhizobiome samples across pH categories (all samples used), also assessed in the same manner as in part C. Statistical significance is indicated as follows: ns (grey) = p > 0.05, * (orange) p < 0.05, ** (red) p < 0.01, *** (brown) p < 0.001.

Our results suggested that local soil communities were significantly influenced by soil pH (ANOVA: DC_LS_ = 7.00 ± 0.74, AC_LS_ = 7.27 ± 0.71, WC_LS_ = 6.17 ± 0.64; Kruskal-Wallis: DC_LS_ vs. WC_LS_ (p = 5.40 × 10^-8^), the DC_LS_ vs. AC_LS_ (p = 0.02), and the WC_LS_ vs. AC_LS_ (p = 1.60 × 10^-5^), whereas in the rhizobiome, root exudates from the host plant could be acting as buffers to maintain pH homeostasis, thus mitigating these differences^95^. Although, in the rhizobiome, the only significant difference from our PERMANOVA pairwise analysis was observed between the DC_RB_ and WC_RB_, (p = 0.003), suggesting that increased rainfall in the wetter regions could have led to the leaching of basic cations from the soil, resulting in a lower pH; while soils with limited rainfall retained these cations, making them more alkaline^96^.

Our findings further indicated that the ratio (R:pH) between rainfall (R) and soil pH was significant in the local soil and rhizobiome across the cluster categories (p = 0.001). The rainfall and pH ratio were significant for all local soil categories (p = 0.003) for all interactions (Supplementary Table S5). However, for the rhizobiome, only the interaction between the DC_RB_ and WC_RB_ was significant (p = 0.003). The relationships intertwining soil pH, rainfall, and bacterial diversity are complicated. While pH and precipitation emerged as primary drivers of change in bacterial diversity^97,98^, the extent to which these two variables directly influenced bacterial community composition remains unclear. However, common trends regarding pH and precipitation impacts on specific microbial alpha diversity indices have been well documented^89,90,98,99^, leading us to investigate these relationships.

We observed that, across the 25 locations, alpha diversity metrics (Supplementary Table S6) were significantly tied to locations (ANOVA: (LS: S_obs_ p = 1.06 × 10^-8^, Faith PD p = 4.91 × 10^-10^, and H’ p = 7.38 × 10^-10^) and (RB: S_obs_ p = 6.56 × 10^-6^, Faith p = 4.54 × 10^-8^, and H’ p = 1.98 × 10^-5^); Figure 2B). On the other hand, while local soil alpha diversity metrics significantly differed across the local soil cluster categories (ANOVA: (LS: S_obs_ p = 6.02 × 10^-5^, Faith PD = 1.55 × 10^-3^, and H’ p = 8.75 × 10^-6^)); there was no significance between the rhizobiome samples possibly due to shielding from the host (Supplementary Table S5).

We examined how pH and precipitation gradients influenced the organization of our cluster categories in terms of microbial diversity and abundance (Supplementary Figure S1). We found that precipitation depth significantly affected all alpha diversity metrics across (ANOVA: LS and RB (S_obs_ p = 0.05, Faith PD p = 3.82 * 10^-3^, H’ p = 7.17 * 10^-3^) and between the sample types ((LS vs. RB) (p < 2 * 10^-16^)). Similar results were observed in our linear regression analysis comparing the alpha diversity metrics of rhizobiomes, ordered by varying levels of rainfall (Supplementary Figure S2).

Pairwise tests further revealed that the most dramatic differences in diversity were observed between the following precipitation (as 60 day-maximum rainfall) categories across sample types: (H’: (LS: p-adj = 2.21 * 10^-3^ (9 to pH 9.5 vs. 12.5 to pH 13), RB: p-adj = 3.21 * 10^-3^ (6.5 to 7 vs. 15 to 15.5)); Sobs: (LS: p-adj = 2.05 * 10^-3^ (9 to 9.5 vs. 12.5 to 13), RB: p-adj = 3.99 * 10^-3^ (6.5 to 7 vs. 16.5 to 17)); and Faith PD: (LS: p-adj = 2.21 * 10^-3^ (13 to 13.5 vs. 16.5 to 17), RB: p-adj = 6.05 * 10^-4^ (14 to 14.5 vs. 16.5 to 17)) (Figure 2C; Supplementary Table S7). While the alpha diversity metrics in our study were more stable across categories of increased rainfall^100^, the local soil revealed the most substantial differences in alpha diversity, mainly among moderate precipitation categories. At the same time, the rhizobiome exhibited more variable changes, particularly between the extreme precipitation groups. This suggests that soil microbial communities were more responsive to subtle deviations in rainfall within our “ambiguous buffer zone” separating arid and humid areas^100,101^. However, because of the mitigating effects of the host, the rhizobiome appeared to be more resistant to intermediate precipitation fluctuations, only showing differences among precipitation extremes^102^.

Linear regression analysis revealed significant changes of relative abundance across the precipitation gradient for seven phyla: *Crenarchaeota*, *Desulfobacterota*, *MBNT15*, *Planctomycetota*, *RCP2-54*, *Verrucomicrobiota*, and *WPS-2* with p-values of p = 4.00 * 10^-2^, p = 1.00 * 10^-2^, p = 9.41 * 10^-3^, p = 2.11 * 10^-2^, p = 1.09 * 10^-2^, p = 3.46 * 10^-4^, and p = 9.64 * 10^-4^ respectively (Supplementary Figure S3). All except *Crenarchaeota* (slope = -1.22 * 10^-4^; R^2^ = 0.306) increased in relative abundance with greater precipitation depth. We hypothesize that the decrease in *Crenarchaeota* might be due to their roles as ammonia-oxidizing archaea (AOA), which might not be in high demand or feasible under waterlogged conditions^103^. In contrast, no significant changes in alpha diversity (ANOVA: (S_obs_ p = 0.73,

Faith PD = 0.59, H’ p = 0.49)) or bacterial relative abundance were found across the pH gradient; with significant changes only observed when comparing sample types (p < 2 * 10^-16^) and categorized cluster regarding Faith PD (p = 2.08 * 10^-3^) (Figure 2D). However, two-way ANOVA revealed significance regarding the relationship between Shannon diversity and pH*cluster category (p = 1.54 * 10^-2^). As with rainfall we conducted a linear regression analysis to compare the alpha diversity metrics of rhizobiomes and local soils, organized by decreasing pH (Supplementary Figure S4).

Pairwise analysis indicated no significant differences in alpha diversity metrics among rhizobiome categories and only minimal differences across local soil pH categories, suggesting that while community composition may be influenced, overall richness remains relatively stable (Supplementary Table S7). The minimal variation in microbial diversity across pH subdivisions suggests that legacy effects of the host plant variety may exert a stronger influence on current plant-microbe associations^10,104,105^. This interpretation is further supported by findings from our previous reciprocal garden experiment^19^, in which host wet varieties exhibited reduced richness (S_Obs_) in dry soil compared to ambiguous and dry host varieties grown under the same conditions.

Collaborating with our previous findings that precipitation is the most critical factor in driving differences in host-associated rhizobiome composition under “abiotic extremes,” led us to ask the following question: How does the interaction between precipitation, sample type, and sample category assignment influence the distribution of ASVs among our samples?

### Prokaryotic community composition revealed patterns that differentiated our clustering categories

We observed that the local soil and rhizobiome samples had similar microbial community composition trends across and within our cluster categories (Figure 3A). Interestingly, we observed that the dry-categorized samples had community compositions more similar to each other than the wet-categorized samples, suggesting that increased abiotic stress might have a filtering effect on the microbial community structures^106,107^.

**Figure 3:**
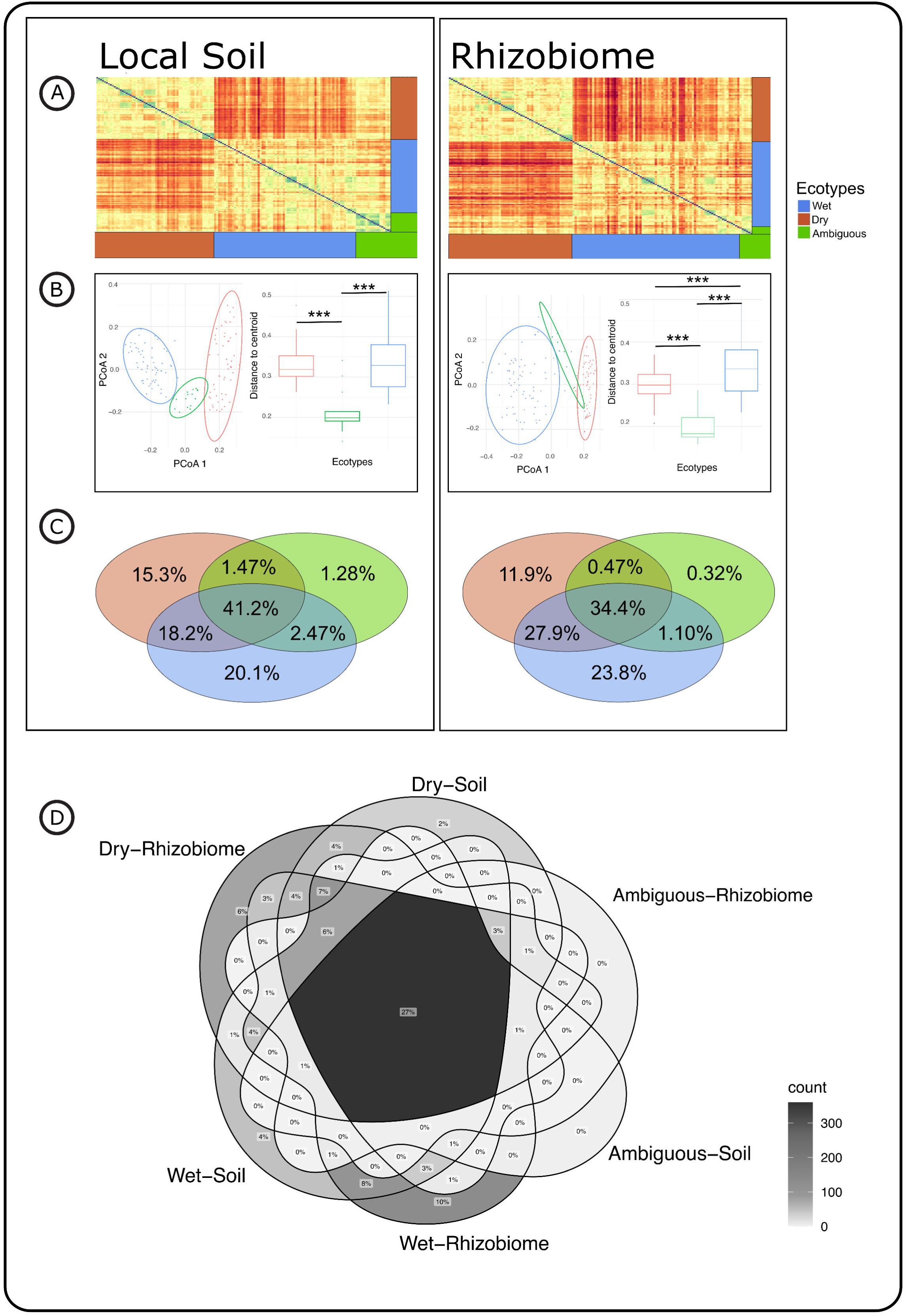
Beta diversity patterns and ASV distribution across cluster categories and sample types. (A) Bray–Curtis dissimilarity heatmaps illustrating large-scale community composition trends in (A) local soil and (B) rhizobiome samples, clustered by their cluster assignments (dry and wet). (B) PCoA plots of beta dispersion, showing community variability within cluster category (dry and wet) for each sample type. The accompanying box plots display the distance of each sample to the group centroid. Statistical significance is defined as follows: ns = p > 0.05, * p < 0.05, ** p < 0.01, *** p < 0.001. (C) Venn diagrams depicting the distribution of shared and unique ASVs across dry and wet cluster categories for local soil and rhizobiome samples. (D) Combined Venn diagram comparing ASV overlap between local soil and rhizobiome sample types across dry and wet cluster categories.

We found that beta diversity dispersion significantly differed among clustering categories for both local soil and rhizobiome communities (PERMANOVA, p = 0.001; F-statistic LS: 28.4, RB: 24.9), with BetaDisper distances to the median exhibiting inconsistent levels of within-group variability (DC_LS_ = 0.33, DC_RB_ = 0.30; AC_LS_ = 0.21, AC_RB_ = 0.20; WC_LS_ = 0.33, WC_RB_ = 0.34). Furthermore pairwise comparisons showed significant differences in betadispersion between all of the categories (DC_LS_ vs. MC_LS_: p = 1.57 × 10^-13^, AC_LS_ vs. WC_LS_: p = 1.35 × 10^-9^; DC_RB_ vs. MC_RB_: p = 1.75 × 10^-7^, AC_RB_ vs. WC_RB_: p = 4.79 × 10^-6^, and DC_RB_ vs. WC_RB_: p = 1.00 × 10^-3^), except for one interaction (DC_LS_ vs. WC_LS_: p = 0.91). These results support our previous findings that the microbial communities within each cluster can vary significantly, with some cluster categories exhibiting more pronounced heterogeneity (dry and wet) than others ambiguous ^19^.

Although the dry and the wet cluster categories did not differ significantly in terms of ASV composition comparing sample types, the following scenarios prompted us to investigate differences in cluster categorized microbial composition: I) We noticed that the dry rhizobiome samples are more similar to each other than dry local soil; II) additionally, the dry rhizobiome composition is very different from the wet rhizobiome (Figure 3B). While the shared ASVs across all cluster categories (41.2% LS and 34.4% RB) represented the highest relative abundance, the wet-categorized cluster had the second-highest number of unique ASVs (20.1% LS and 23.8% RB), followed by the dry-categorized cluster (15.3% LS and 11.9% RB) (Figure 3C). Across the sample types (LS and RB), we observed that 26.5% of ASVs were shared among all categories and samples (Figure 3D, Supplementary Table S8). The wet-categorized cluster had more unique ASVs than the dry-categorized cluster. We deduce that this difference could be linked to various microbial legacy effects and adaptation to abiotic stress^108,109^. In this context, the dry-categorized cluster might prefer specialist microbes over generalists, reducing the overall number of unique ASVs^110,111^. Correspondingly, the small number of unique ASVs between clusters and sample types might result from phylogenetic divergence and variations in niche space, where microbes adapted to specific precipitation pressures may not suit contrasting environments^112^. We further confirmed that there were significant differences in unique ASVs across cluster categories (Chi-squared, p = 2.2 × 10^-16^), and between wet and dry categories (p = 0.05), demonstrating that differences in unique ASVs were primarily due to cluster category, rather than sample type (p = 0.557). Our results suggest that complex environmental and climate factors have a more profound impact on shaping the number of unique ASVs than the localized effects of individual plant hosts.

### Stochastic processes influenced community composition more in the local soil than in the rhizobiome in arid locations

Both local soil and rhizobiome samples showed high beta dispersion distances across all of our dry and wet locations and cluster categories, common among grassland microbiomes (Figure 4A, 4B; Supplementary Table S9)^113^. We observed notable differences between the dry cluster categorized local soil and rhizobiome communities, in which 8 out of 10 comparisons showed significant differences in beta dispersion (Dry: TX3: p = 3.43 × 10^-5^, NE1: p = 5.41 × 10^-6^, OK1: p = 1.45 × 10^-11^, KS2: p = 3.08 × 10^-2^, ND1: p = 1.20 × 10^-3^, MT1: p = 6.02 × 10^-3^, SD1: p = 5.06 × 10^-3^, NM1: p = 1.52 × 10^-3^, and CO1: p = 8.69 × 10^-4^), whereas only one comparison for wet-categorized clusters was significant (Wet: MI1: p = 3.02 × 10^-2^).

**Figure 4:**
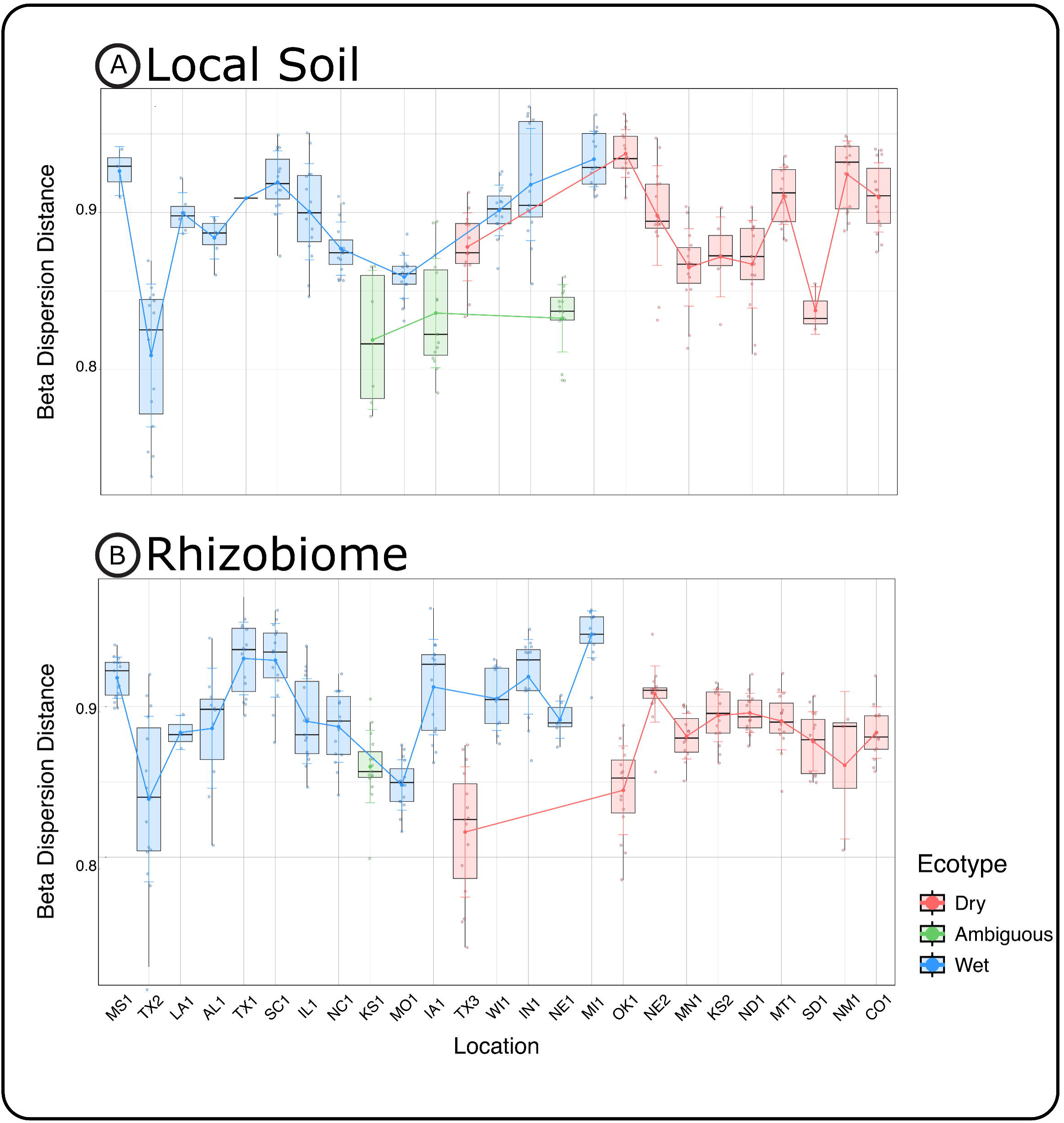
Within-site community variability across local soil and rhizobiome samples. (A) Beta dispersion analysis of local soil samples, assessing within-site variability across locations. Sample points are colored by their cluster category, with lines representing the mean distance to the centroid for each category and shaded bars indicating standard deviation. Sites are arranged in order of decreasing precipitation. (B) Beta dispersion analysis of rhizobiome samples, with sample coloring and site arrangement as described in part A. Statistical comparisons between local soil and rhizobiome samples at each site indicate significant differences in beta dispersion, suggesting variation in community stochasticity across sample types. Statistical significance is denoted as follows: ns = p > 0.05, * p < 0.05, ** p < 0.01, *** p < 0.001.

Notably, among the two driest locations, CO1 and NM1, the local soil had significantly higher beta dispersion than their rhizobiome communities, a trend unique to the dry-categorized clusters. We hypothesize that dispersal limitation and host influence on microhabitat stability and selection might drive the observed differences in stochasticity among various sample types in the driest locations. This could result from dry clusters reducing dispersal effects by fostering specialized microbial communities shaped by abiotic stress, leading to a more stable rhizobiome community structure than the local soil^114^. With the dry-associated cluster category rhizobiome demonstrating less stochasticity, we next asked what microbial populations in the rhizobiome could contribute to the similarity and stability associated with the dry variety plant hosts.

### Oligotyping analysis reveals contrasting variant compositions across cluster categories and sample types for six prokaryotic families

To analyze fine-scale ecological patterns in local soil and rhizobiome samples exclusively across wet and dry cluster categories, we conducted an unbiased taxonomic oligotype assessment for all our ASVs. Our clustering for oligotyping analysis mirrored the clustering of our ASVs. This suggests that the impact of rare oligotypes within each taxon from each location was limited, while precipitation and ecological pressures drove common trends between ASVs and their variants at each location (Supplementary Figure S5, Supplementary Table S10).

For both sample types, the permutation test for homogeneity revealed significant differences in betadispersion between the cluster categories for each sample type (LS and RB: p = 0.001, LS: F = 10.9, RB: F = 14.2). However, we only observed differences between the dry and wet categorized clusters within the rhizobiome ((LS distance to centroid: 0.24 and 0.26, p = 0.13) and (RB distance to centroid: 0.21 and 0.26, p = 4.25 x 10^-6^) for dry and wet, respectively) (Figure 5A). The distinctions between dry and wet rhizobiomes could originate from more substantial selection pressures and reduced microbial diversity at our drier locations. Furthermore, the contrast between dry local soil and rhizobiome might be more pronounced due to host selection^115^, microhabitat stabilization^116^, and environmental filtering^110^.

**Figure 5:**
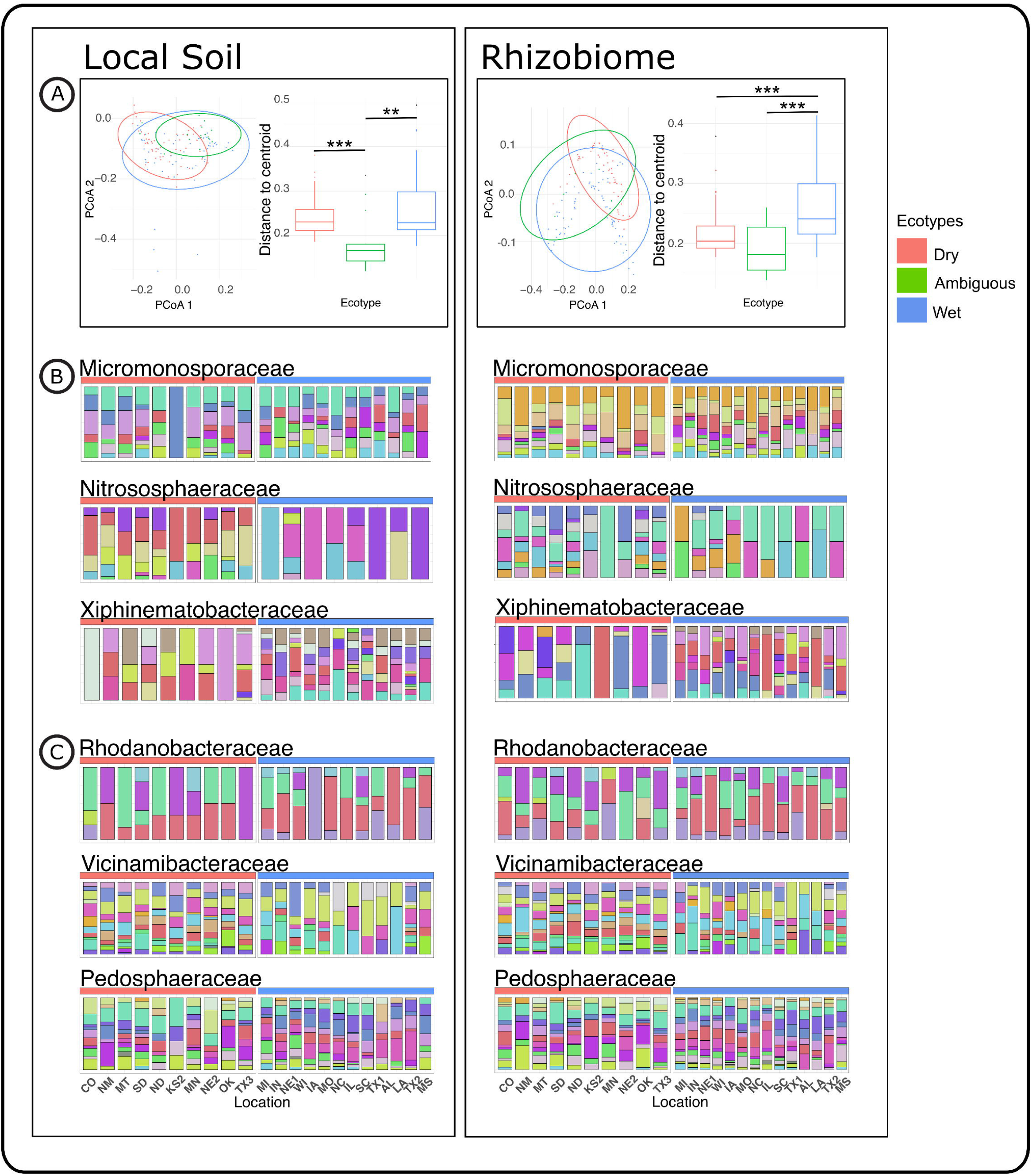
Community variant structure and oligotype composition across cluster category and sample types. (A) PCoA plots of beta dispersion based on taxonomically unbiased oligotyping analysis, illustrating community variability within cluster categories for each sample type. Accompanying box plots display the distance of individual samples to their group centroid, describing within-group variability. Statistical significance is denoted as follows: ns = p > 0.05, * p < 0.05, ** p < 0.01, *** p < 0.001. (B) The top panels represent host-selected communities. Stacked bar plots showing the relative abundance (y-axis) of individual oligotypes within selected prokaryotic families, stratified by sample type and dry versus wet-categorized locations (x-axis). Each color corresponds to a distinct oligotype variant. Details of the oligotypes are in Supplementary Tables 10 and 11. (C) Oligotypes of microbial populations hypothesized to be non-host-selected communities. Stacked bar plots showing the relative abundance (y-axis) of individual oligotypes within selected prokaryotic families, stratified by sample type and dry versus wet-categorized locations (x-axis). Each color corresponds to a distinct oligotype variant. Details of the oligotypes are in Supplementary Tables 10 and 11.

Given the differences between the dry and wet categorized clusters and across the two sample types, we aimed to evaluate the oligotypic patterns within specific ASV families to examine the abundance of particular community oligotypes among closely related species of microorganisms (Figure 5B and 5C; Supplementary Table S11). Taxonomic analysis revealed distinct results across the categorized clusters and sample types among the *Micromonosporaceae, Nitrosphaeraceae*, and *Xiphinematobacteraceae* versus the *Vicinamibacteraceae, Rhodanobacteraceae,* and *Pedosphaeraceae* families. For the *Micromonosporaceae, Nitrosphaeraceae*, and *Xiphinematobacteraceae* families, the highest relatively abundant oligotypes in the local soil differed significantly from those in the rhizobiome, with very few “core” variants between sample types (Figure 5B). The similarities between the dry and wet rhizobiome oligotypes compared to the local soil suggested that host selection might favor specific oligotypes over others^117^, regardless of the location and variant composition of the local soil for members of the *Micromonosporaceae* family. This observed host influence might stem from notably, that many species in the *Micromonosporaceae* family are frequently associated with plant rhizospheres due to their production of plant growth regulators, including auxins and gibberellins^118,119^. Additionally, the *Nitrosphaeraceae* family constitutes ammonia-oxidizing archaea (AOA), which may benefit host nutrient acquisition^120^. The fact that both of these families had similar oligotypic distribution patterns supported the “stress-induced selection” hypothesis, where, under abiotic stress, plants would likely retain the microorganisms within the rhizobiome that benefit the plant^121,122^.

However, in the *Vicinamibacteraceae, Rhodanobacteraceae,* and *Pedosphaeraceae* families, there was greater similarity of oligotypes across the cateorgized clusters and sample types, indicating a more extensive variety of “core” variants (Figure 5C). While more shared oligotypes existed, each location was more distinct than observed in the *Micromonosporaceae* family. We speculated that these differences arose because members of the *Pedosphaeraceae* family are more “free-living” than those associated with the *Micromonosporaceae* family^123^. Because of this minimized host selection, its local soil and rhizobiome oligotype composition were more similar within a location, meaning that the oligotypes found in the rhizobiome most likely originated from the local soil, with little to no host filtering^116,117^.

Despite these apparent distinctions among these families, common trends emerged. Generally, the wet local soil and rhizobiome contained more oligotypes than the dry local soil and rhizobiome locations, reflecting the filtering effect from both the environment and plant host^124,125^. Additionally, the rhizobiome consistently had more oligotypes than the local soil, suggesting that the plant host could support a higher diversity of microorganisms within a family due to increased nutrients and reduced fluctuating stress associated with the environment. Additionally, we theorized that microhabitat stability and the increased number of features present for the rhizobiome in both families (*Micromonosporaceae*: LS: n = 329 and RB: n = 587; *Xiphinematobacteraceae:* LS: n = 251 and RB: n = 305; *Nitrosphaeraceae:* LS: n = 196 and RB: n = 205; *Vicinamibacteraceae:* LS: n = 758 and RB: n = 932; *Rhodanobacteraceae:* LS: n = 136 and RB: n = 318; *Pedosphaeraceae*: LS: n = 755 and RB: n = 1,083) might have contributed to the rhizobiome samples having more variants regardless of cluster assignment^126^. Considering the oligotypic analysis here, we suggest that as the abiotic stress of the environment increases, the host influence on the rhizobiome community exerts a greater pressure.

## Conclusion

Precipitation significantly shapes the compositions of microbiome communities^127^. However, discrepancies between precipitation and the strength of host interaction can lead to varying impacts on community structure, some of which can be exacerbated under increased abiotic stress^128,129^. Here, we demonstrated that regional precipitation profoundly affected and categorized the microbiomes of *A. gerardii* into two distinct cluster categories (i.e., dry, wet) and one ambiguous cluster category that comprised the intersection between the dry and the wet clusters. Notably, the geographic span of our categorized clusters crossed distinguishable areas of the 100^th^ meridian, known as North America’s “arid-humid divide.” We speculate that our categorized clusters’ composition and geographic span might be linked to the *A. gerardii* host varieties’ diversity and legacy effects^9,12,130^.

Additionally, our findings showed that although pH and precipitation depth had a more pronounced effect on local soil than on rhizobiome communities, alterations in rhizobiome diversity were more pronounced at precipitation extremes. We surmise that these differences might arise from host variety (ecotypic) effects through microhabitat stabilization. By stratifying our samples by type and respective clusters, we further identified large-scale trends in microbial community composition, particularly in the differences between the dry and wet categorized clusters. In this study, we observed significant variations in the relative abundances of specific phyla within sample-type clusters. This suggests specialized roles of specific phyla responding to precipitation-related stress and underscores potential precipitation-community structure relationships.

Furthermore, for the first time, we showed notable differences in rhizobiome and local soil microbial diversity with reduced precipitation, revealing possible links between abiotic stress and microbial community assembly strategies (stochastic vs. deterministic). We further demonstrated that regional precipitation was a critical factor in rhizobiome assembly, with the rhizosphere communities in the aridest locations exhibiting more predicted forms of randomness than the local soil, implying plant host-associated biotic filtering under extreme abiotic stress (Figure 6). Our findings highlight the shift toward reduced stochasticity in the microbial communities under intensified abiotic stress. This underscores the pivotal role of regional precipitation in shaping plant-associated rhizobiomes and suggests that plant hosts might rely heavily on biotic filtering mechanisms to mitigate extreme environmental challenges.

**Figure 6:**
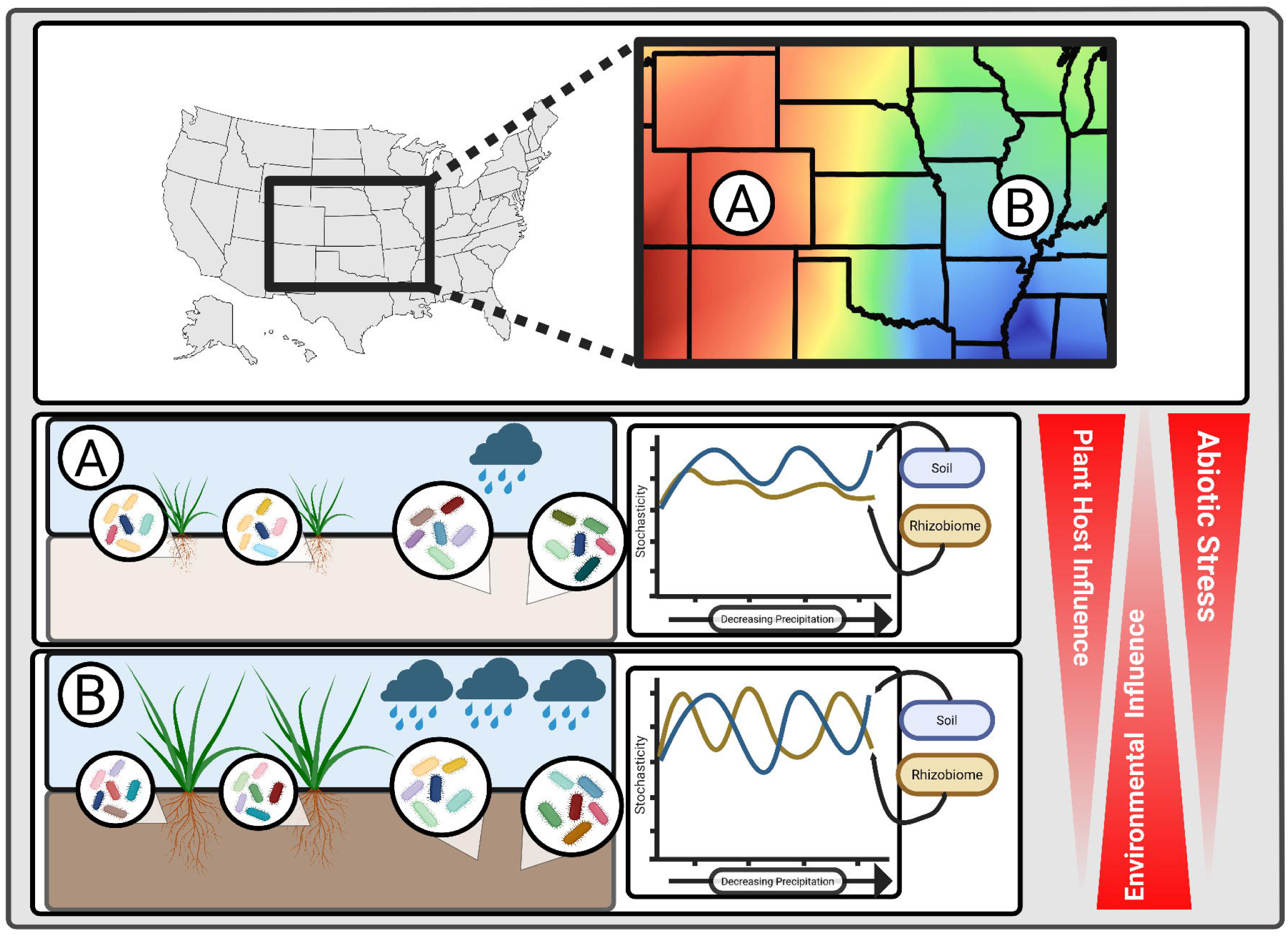
Conceptual model summarizing the influence of precipitation-driven abiotic stress on soil and rhizobiome community assembly. The top panel illustrates a subdivision of the United States precipitation gradient, highlighting two locations, A and B, to describe two scenarios based on our findings. The second panel (Location A) illustrates a high-stress environment with limited precipitation. In this scenario, plant host selection has a strong influence on rhizobiome community assembly, leading to divergence from the surrounding soil community and the enrichment of specialist taxa. This shift reflects a reduction in environmental influence and an increase in host-mediated filtering under abiotic stress. Here, the stochasticity of the rhizobiome becomes more uniform, with more similar plant-associated rhizobiome compositions, as site aridity increases while the local soil remains highly variable. In contrast, the bottom panel (Location B) depicts a low-stress environment characterized by high precipitation. Under these conditions, environmental influences dominate, while the impact of plant hosts is minimal. Consequently, local soil and rhizobiome communities exhibit high variability. Generalist taxa dominate both communities. Here, the stochasticity of the local soil and rhizobiome remains high, showing a highly variable plant-associated rhizobiome, and precipitation has a minimal effect.

## Acknowledgments

We thank everyone involved in sampling and data acquisition; this research would not have been possible without your contributions. We also thank Tanner Riche and Qinghong Ran for their helpful discussions on data analysis. We appreciate the work conducted by Alina Akhunova, Jose Ruiz Ortiz, and Samantha Elledge at the Integrated Genomic Facility at Kansas State University (https://www.k-state.edu/igenomics/). In addition, we thank the Kansas State University Soil Testing Laboratory for helping with the soil property analyses (https://www.agronomy.k-state.edu/outreach-and-services/soil-testing-lab/).

*We acknowledge that this research was conducted on land that is the homeland of many Native Nations, who have long served as stewards of North American tallgrass prairies. We honor their histories, cultures, and traditions and recognize their contributions to the scientific, ecological, and historical knowledge of tallgrass prairie ecosystems.*

## Author Contributions

A.K., J.S., D.C., H.W., M.W., S.B., A.J., L.J., and S.T.M.L. collected and processed samples. B.V., B.A., L.R., R.K., D.C., and D.T. extracted, quantified, and normalized genomic DNA. B.V. and A.K. conducted the bioinformatic analysis of 16S rRNA sequencing. B.V. carried out the statistical analyses. B.V. and S.T.M.L. authored the manuscript, developed figures, and prepared all supplementary files. A.J. and L.J. assisted with revisions of the manuscript and figures. S.T.M.L., A.J., L.J., M.W. and S.B. secured funding for this study. All authors reviewed, edited, and endorsed the last version of this manuscript.

## Author’s Information

Sonny T.M. Lee:

X: @sonnytmlee

Bluesky: @sonnytmlee.bsky.social

LinkedIn: https://www.linkedin.com/in/sonny-tm-lee-a25b9120 ResearchGate: https://www.researchgate.net/profile/Sonny-Lee-3

Brooke Vogt:

X: @VogtBrookeM

Bluesky: @brookevogt.bsky.social

LinkedIn: www.linkedin.com/in/brooke-vogt-5a351035b

ResearchGate: https://www.researchgate.net/profile/Brooke-Vogt

## Supplementary Figures

**Supplementary Figure S1:**
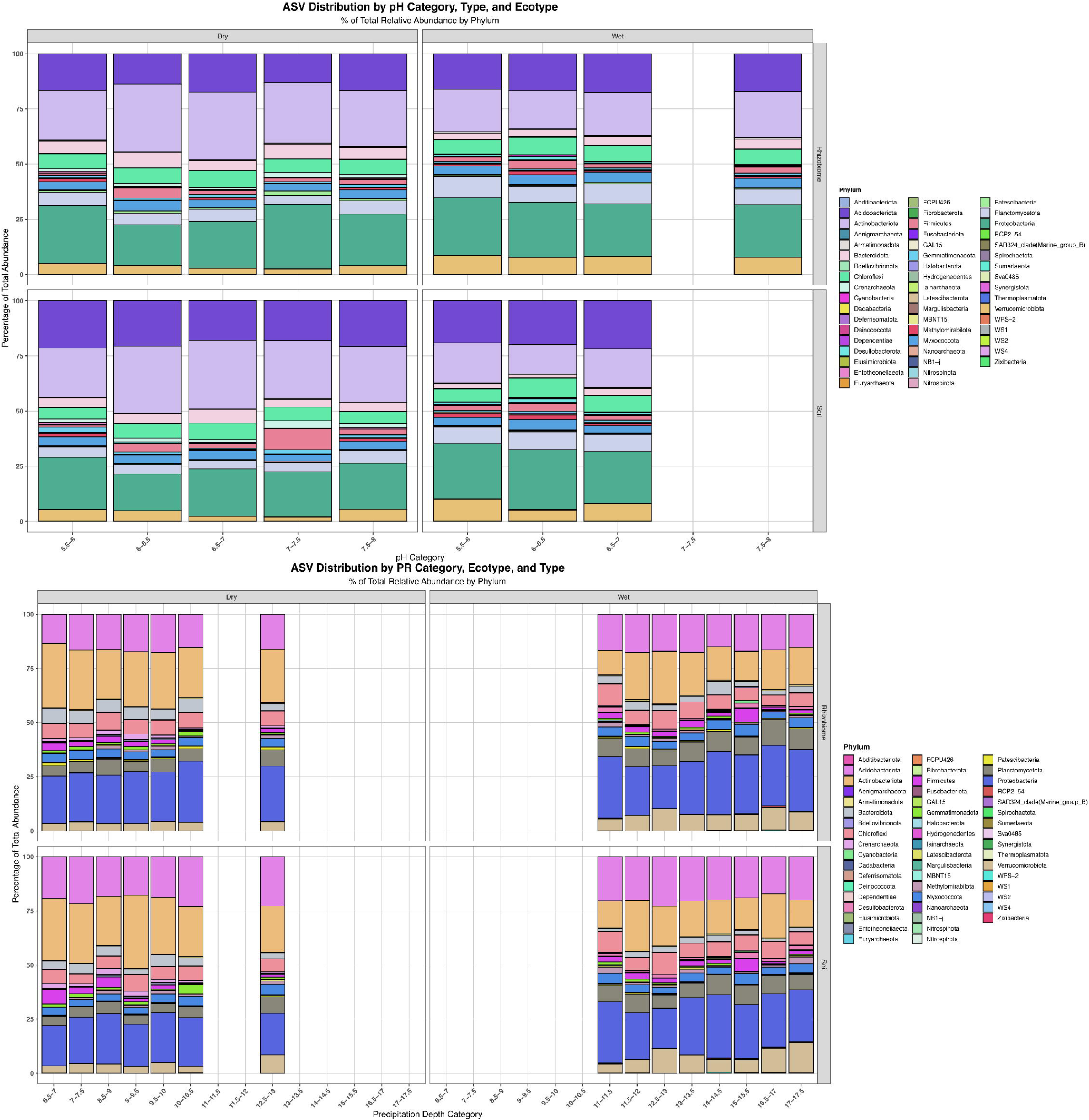
Microbial community composition across climate and soil property gradients. Stacked bar plots showing the relative abundance of ASVs classified at the phylum level, stratified by cluster category, environmental pH categories (top), and precipitation (bottom) gradients. This visualization highlights compositional shifts in microbial communities along gradients of climate and soil properties.

**Supplementary Figure S2:**
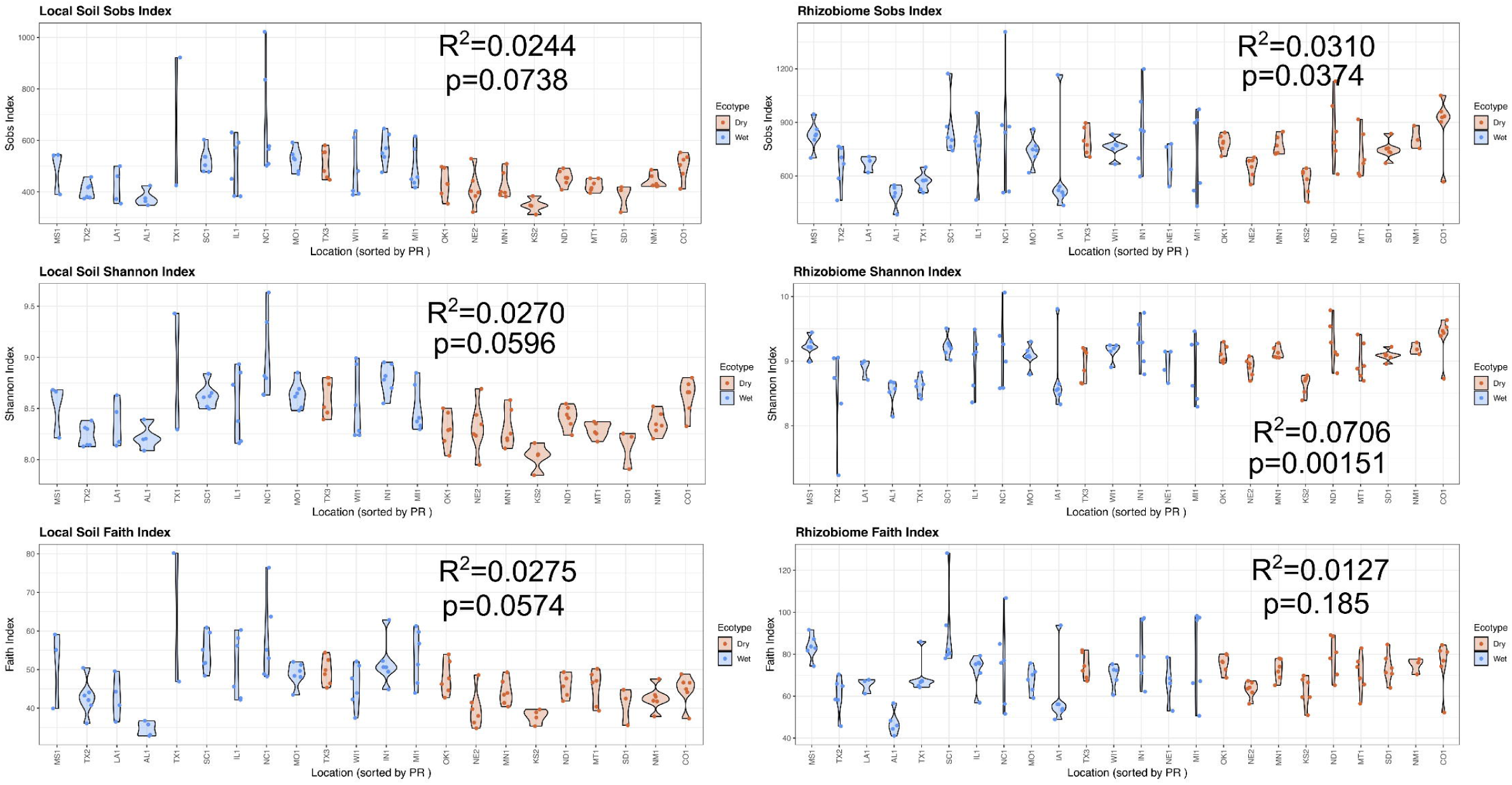
Precipitation linear regression analysis on alpha diversity. Linear regression analysis was conducted to evaluate the relationship between precipitation and alpha diversity metrics—including S_obs_, H’, and Faith PD—in local soil (left panels) and rhizobiome (right panels) communities. Samples were arranged along a gradient of decreasing precipitation and visualized using violin plots. The plot color denotes the location’s cluster category (dry and wet). Displayed in each of the plots are the p and R^2^ values from the regression analysis.

**Supplementary Figure S3:**
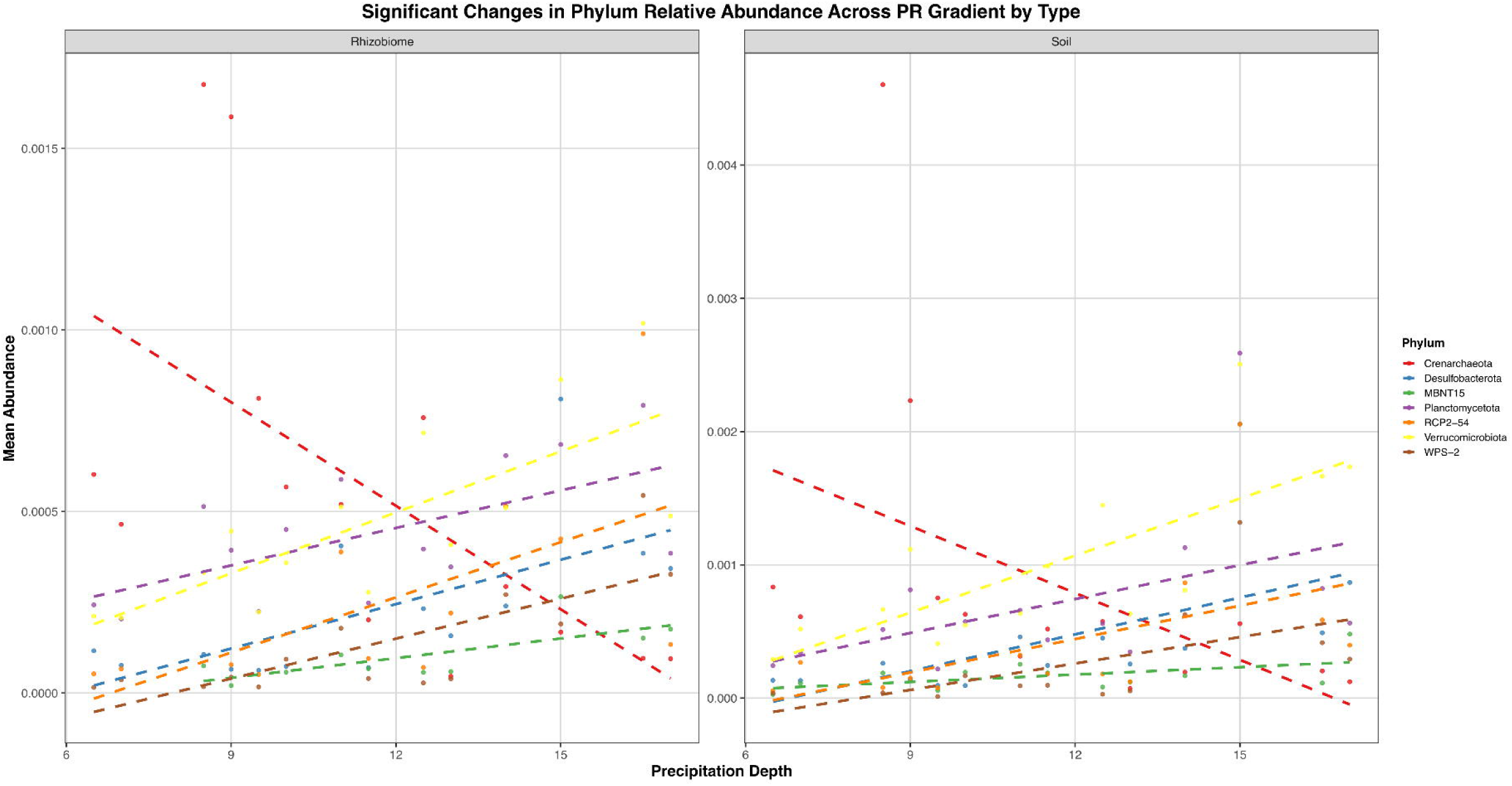
Linear regression analysis of phylum-level microbial abundance across categories and sample types with increasing precipitation depth. Linear models assessed phyla whose relative abundances significantly correlated with precipitation across different categories and sample types (local soil and rhizobiome). Only phyla exhibiting significant relationships with precipitation (p < 0.05) are shown, highlighting taxa potentially responsive to moisture variability along the precipitation gradient.

**Supplementary Figure S4:**
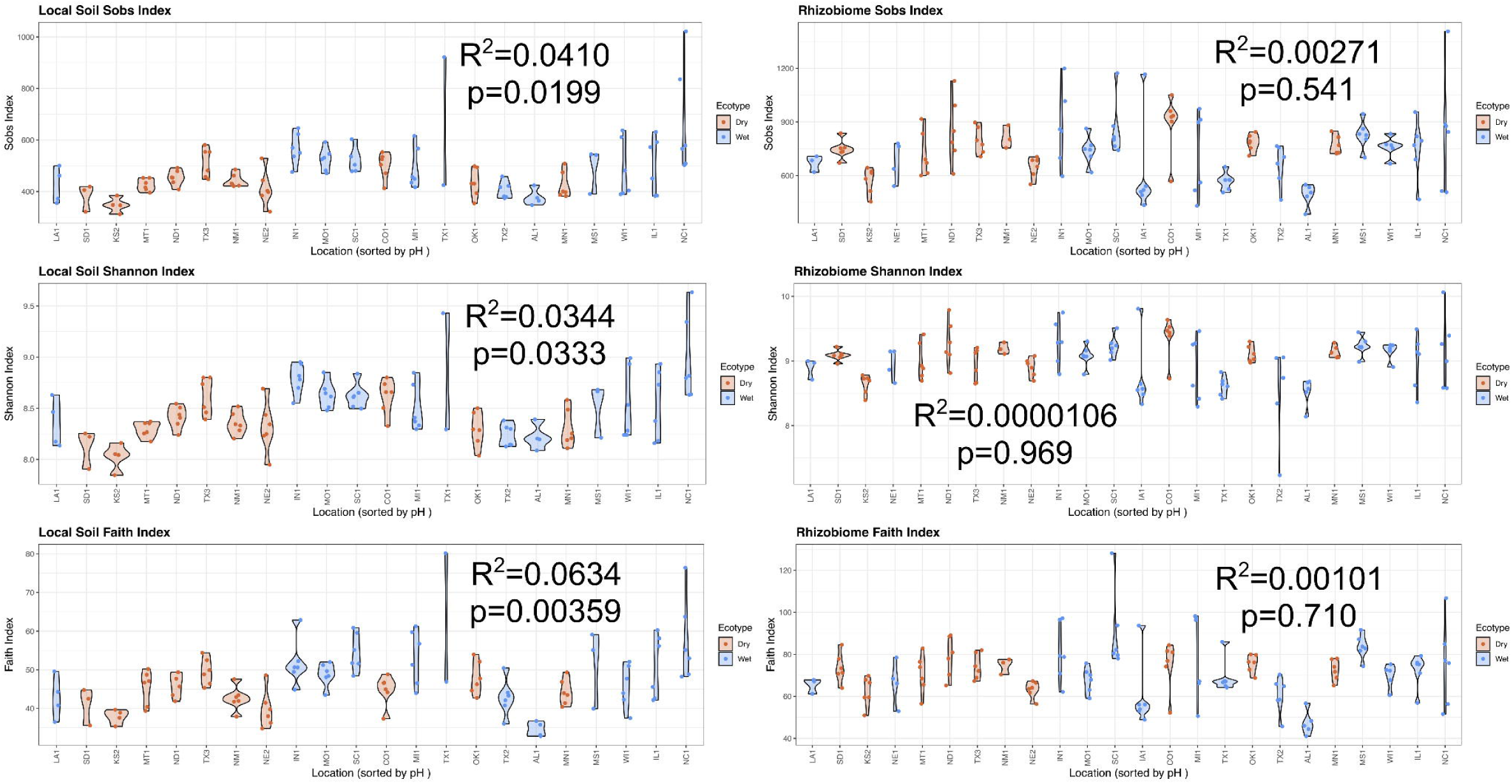
pH linear regression analysis on alpha diversity. Linear regression analysis was conducted to evaluate the relationship between pH and alpha diversity metrics—including S_obs_, H’, and Faith PD—in local soil (left panels) and rhizobiome (right panels) communities. Samples were arranged along a gradient of decreasing pH and visualized using violin plots. The plot color denotes the location’s cluster category (dry and wet). Displayed in each of the plots are the p and R^2^ values from the regression analysis.

**Supplementary Figure S5:**
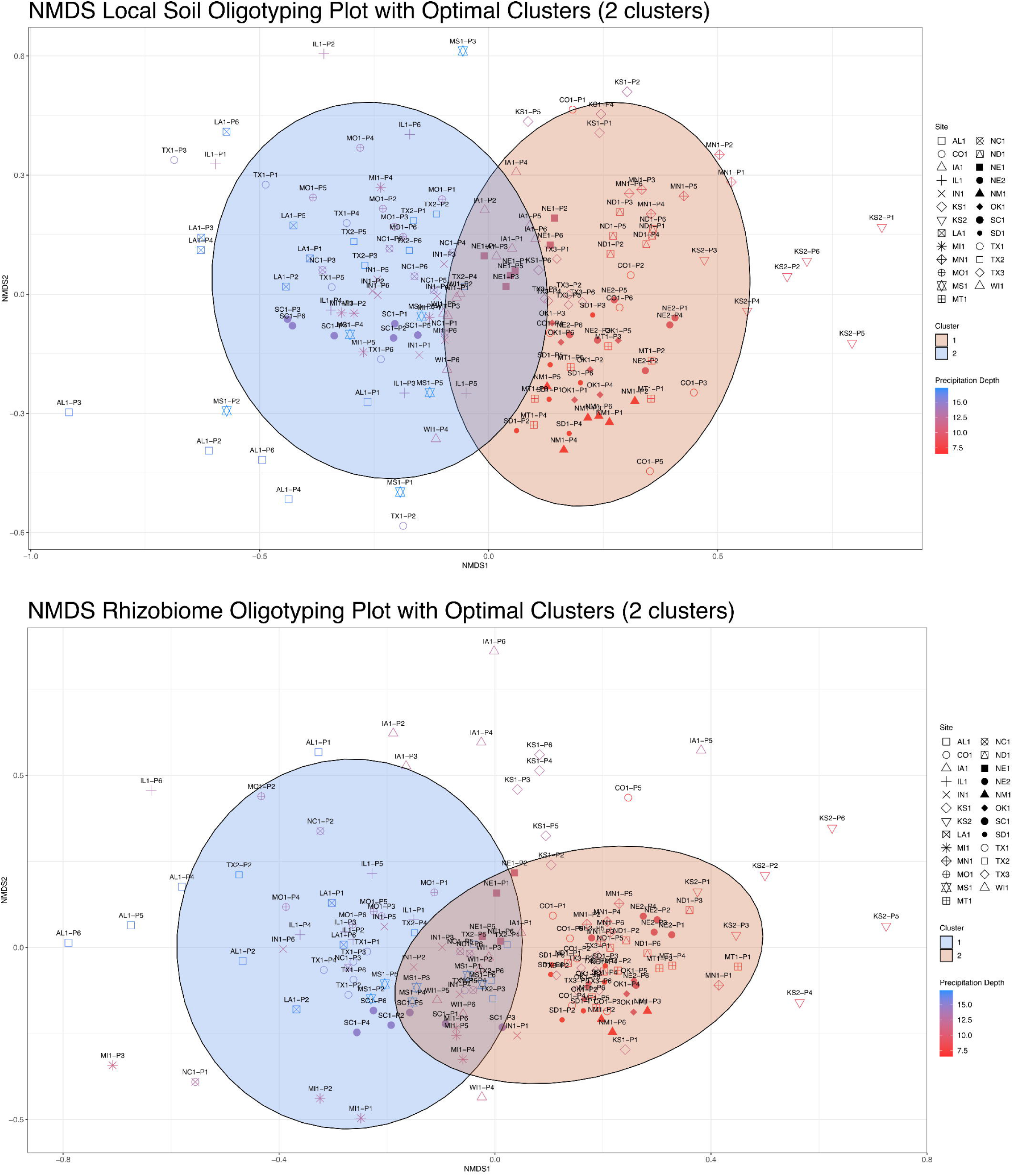
Taxonomically unbiased oligotype clustering. NMDS taxonomically unbiased oligotype clustering of local soil (top) and rhizobiome (bottom) variants within each sample with k-means clustering and optimized silhouette scores showing distinct groupings with 95% confidence ellipses. Each point represents a sample, colored according to NOAA’s 60-day maximum precipitation data at each site.

## Supplementary Tables

Supplementary Table S1: Site Locations with Precipitation Metrics

Supplementary Table S2: QIIME Local Soil and Rhizobiome Sequences Before and After Quality Trimming

Supplementary Table S3: ASV Assignments for the Local Soil and Rhizobiome Samples

Supplementary Table S4: Soil Chemistry

Supplementary Table S5: Soil Properties and Alpha Diversity Statistical Analyses

Supplementary Table S6: Alpha Diversity for the Local Soil and Rhizobiome Samples

Supplementary Table S7: Precipitation and pH Gradient t-Test Results

Supplementary Table S8: ASV Distribution by Sample Type and Cluster Category

Supplementary Table S9: Beta Dispersion Distance for Local Soil and Rhizobiome Samples within a Location

Supplementary Table S10: Unbiased Taxonomic Oligotyping Results

Supplementary Table S11: Oligotyping Results for Individual Families

